# *In vivo* discovery of RNA proximal proteins in human cells via proximity-dependent biotinylation

**DOI:** 10.1101/2020.02.28.970442

**Authors:** Xianzhi Lin, Marcos A. S. Fonseca, Rosario I. Corona, Kate Lawrenson

**Affiliations:** Women’s Cancer Research Program at Samuel Oschin Comprehensive Cancer Institute, Cedars-Sinai Medical Center, Los Angeles, CA, USA; Division of Gynecologic Oncology, Department of Obstetrics and Gynecology, Cedars-Sinai Medical Center, Los Angeles, CA, USA; Center for Bioinformatics and Functional Genomics, Samuel Oschin Comprehensive Cancer Institute, Cedars-Sinai Medical Center, Los Angeles, CA, USA

**Author notes:** Correspondence (X.L.) or (K.L.).

**Keywords:** Type VI CRISPR-Cas (Cas13), Engineered soybean ascorbate peroxidase APEX2, Proximity-dependent biotinylation, RNA proximity labeling (RPL), RNA proximal proteins (RPPs), RNA-centric method, RNA-protein interactions, poly(A) tail, RNA binding proteins (RBPs), *U1* snRNA interactors

## Abstract

RNA molecules function as messengers or noncoding adaptor molecules, structural components, and regulators of genome organization and gene expression. Their roles and regulation are mediated by other molecules they interact with, especially RNA binding proteins (RBPs). Here we report RNA proximity labeling (RPL), an RNA-centric method based on fusion of an endonuclease-deficient Type VI CRISPR-Cas protein (dCas13b) and engineered ascorbate peroxidase (APEX2) to discover *in vivo* target RNA proximal proteins (RPPs) through proximity-based biotinylation. *U1* RPPs enriched by proximity-based biotinylation included both *U1* snRNA canonical and noncanonical functions-related proteins. In addition, profiling of poly(A) tail proximal proteins uncovered expected categories of RBPs for poly(A) tails and also provided novel evidence for poly(A)+ RNA 5’-3’ proximity and expanded subcellular localizations. Our results suggest that RPL is a rapid approach for identifying both interacting and neighboring proteins associated with target RNA molecules in their native cellular contexts.

## Introduction

RNA molecules include both messengers encoding proteins (mRNAs) and noncoding RNAs (ncRNAs) such as adaptor tRNAs and regulatory long noncoding RNAs (lncRNAs). Around 2% of the human genome encodes mRNAs (International Human Genome Sequencing Consortium, 2001; Venter et al., 2001), while the majority is pervasively transcribed into ncRNAs (Berretta and Morillon, 2009; Djebali et al., 2012), including lncRNAs that are widely considered as a large family of potential regulators (Batista and Chang, 2013; Iyer et al., 2015; Yang et al., 2014). However, only a small number of lncRNAs have been functionally and mechanistically studied and most remain uncharacterized (Kopp and Mendell, 2018).

The functions and regulation of RNA transcripts are mediated by other molecules they associate with, particularly RNA binding proteins (RBPs) that govern many critical RNA activities (Dreyfuss et al., 2002; Glisovic et al., 2008; Hentze et al., 2018; Lunde et al., 2007). Discovery of the interacting proteins for a given transcript plays pivotal role in unveiling its function and underlying mechanism. Currently, mechanistic study of lncRNAs is impeded by the shortage of RNA-centric tools and the limitations of existing methods (Ci Chu et al., 2015; Ramanathan et al., 2019). Antisense probe-based ChIRP (C. Chu et al., 2015) or RAP (McHugh et al., 2015) requires crosslinking via chemicals or UV light. However, chemicals such as formaldehyde also crosslink protein-protein interactions, which may lead to false-positive associations (Panhale et al., 2019). Since UV-crosslinking has very low efficiency, antisense probe-based purification methods usually require a large number of cells (∼100-800 million) (Lin et al., 2019; McHugh et al., 2015), which may not be feasible for slow-growing model systems such as primary cell cultures. Moreover, UV-crosslinking can induce RNA alterations like modifications (Wurtmann and Wolin, 2009) that could change binding affinity of RNA to certain RBPs (Bernard et al., 2012) and impair downstream protein analysis (Urdaneta and Beckmann, 2019). An alternative approach, tagging of endogenous RNA requires genetic manipulation and may interfere with endogenous RNA functions (Laprade et al., 2020). Therefore, methods to discover endogenous RNA interacting proteins are needed.

In this study, we developed RPL (RNA proximity labelling) method to identify in vivo target RNA proximal proteins (RPPs) without crosslinking or genetic manipulation. *U1* RPPs recalled *U1* functional relevant proteins, while poly(A) tail RPPs recalled expected categories of RBPs for poly(A) tails providing additional evidence for poly(A)+ RNA 5’-3’ proximity and expanded subcellular localizations.

Results Design and development of RPL, an RNA-centric method for screening RPPs Inspired by the applications of RNA-targeting Type VI CRISPR-Cas systems (Abudayyeh et al., 2017; Cox et al., 2017; Konermann et al., 2018; Yan et al., 2018) and proximity labeling using engineered soybean ascorbate peroxidase (Lam et al., 2015; Rhee et al., 2013) and biotin ligase (Branon et al., 2018; Kim et al., 2016; Roux et al., 2012), we designed RPL, an RNA-centric approach based on a fusion protein of endonuclease-deficient Cas13 (dCas13) and proximity labeling enzyme APEX2 (Figure 1A). The fusion protein is directed to target RNA by a sequence-specific guide RNA (gRNA). In the presence of hydrogen peroxide (H_2_O_2_X APEX2 in the fusion protein oxidizes substrate biotin-phenol (BP) to short-lived biotin-phenoxyl radicals, which covalently react with electron-rich amino acids (like tyrosine) on RPPs within a small radius (Rhee et al., 2013) of the fusion protein (Figure 1A). The biotinylated RPPs, which may include target RNA direct binding proteins, indirect binding proteins, and proximal proteins just present within biotinylating radius, can be readily enriched using streptavidin beads and profiled by liquid chromatography-tandem mass spectrometry (LC-MS/MS) (Figure 1A).

**Figure 1.**
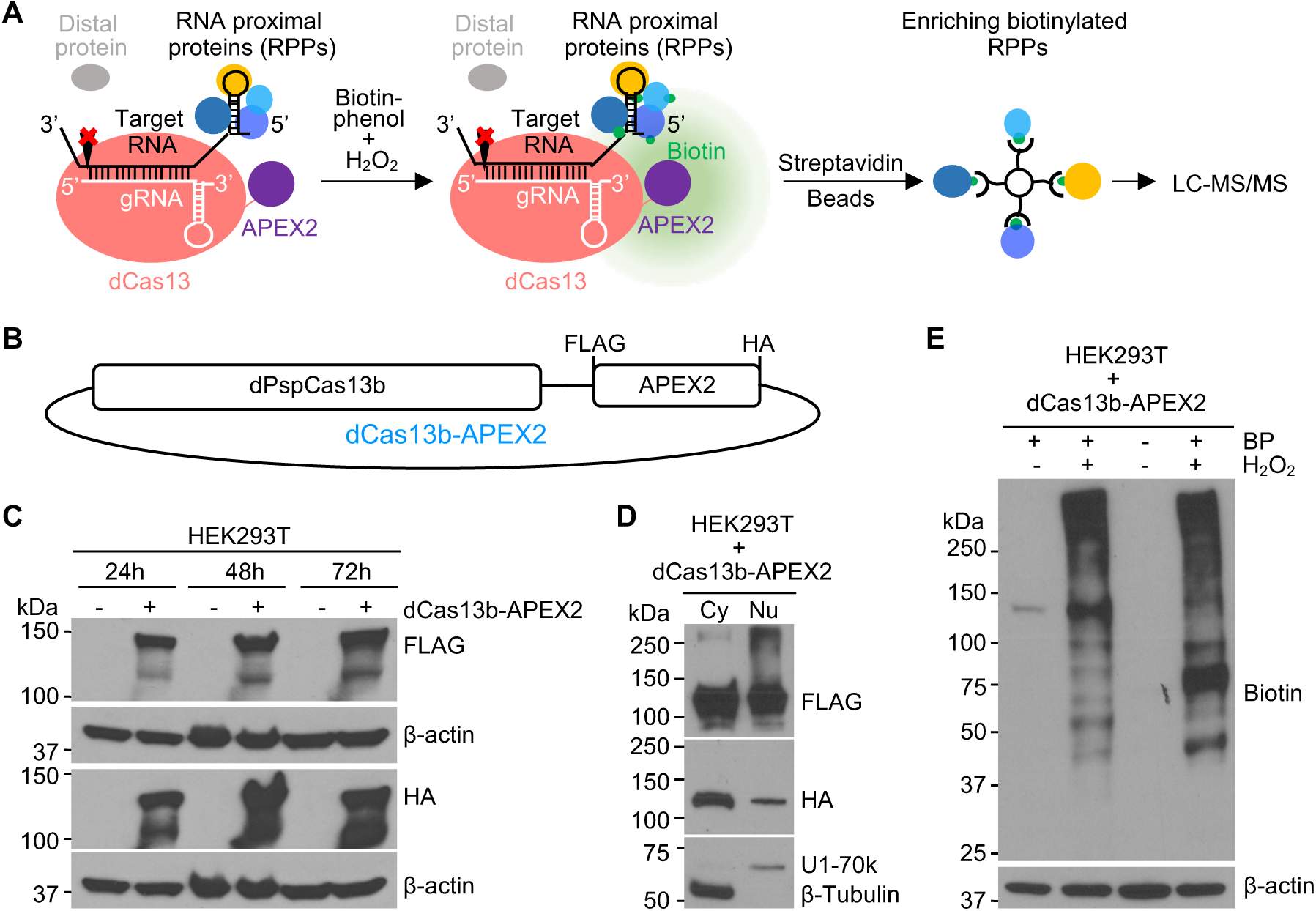
Designing and developing the RPL method. **(A).** Schematic illustration of RPL workflow. A sequence-specific gRNA directs dCas13-APEX2 to target RNA and APEX2 in the fusion protein biotinylates target RNA proximal proteins (RPPs) *in vivo* in the presence of biotin-phenol and H_2_O_2_. Biotinylated RPPs are then enriched using streptavidin beads and analyzed by liquid chromatography-tandem mass spectrometry (LC-MS/MS). **(B).** Diagram of the fusion protein dPspCas13b-FLAG-APEX2-HA (dCas13b-APEX2, or the RPL protein) expression construct. **(C).** Expression validation of the RPL protein by western blot. HEK293T cells transfected with or without the RPL plasmid were harvested 24h-72h post transfection and whole cell lysates were blotted with an anti-FLAG or anti-HA antibody. **(D).** The RPL protein is expressed in both cytoplasm and nucleus. HEK293T cells transfected with the RPL plasmid for 24h were fractionated into cytoplasmic (Cy) and nuclear (Nu) fractions. Fractionation efficiency was evaluated by blotting cytoplasmic protein β-Tubulin and nuclear protein U1-70k. **(E).** Validation of enzymatic activity of APEX2 in the RPL protein. HEK293T cells transfected with the RPL plasmid were treated with different combinations of biotin-phenol (BP) and H_2_O_2_. Whole cell lysates were blotted with anti-biotin antibody, β-actin in **(C)** and **(E)** was used as loading control.

To construct the fusion protein, Cas13b was used for its high efficacy in RNA knockdown with minimal off-target effect (Cox et al., 2017) and high specificity in RNA labeling (Yang et al., 2019). For proximity labeling enzyme, we chose APEX2 for its fast kinetics and high activity (Lam et al., 2015). Catalytically dead Cas13b from Prevotella sp. P5-125 (dPspCas13b) (Cox et al., 2017) was fused to APEX2 with FLAG and HA tags (Figure 1B). The expression of the fusion protein dCas13b-APEX2 (from hereon in called the RPL protein) was confirmed by western blot using an anti-FLAG or anti-HA antibody (Figure 1C). The subcellular localization of the RPL protein was examined when ectopically expressed in HEK293T cells. Efficient separation between cytoplasmic and nuclear fractions was confirmed by blotting for cytoplasmic marker β-Tubulin and nuclear marker U1-70k. The RPL protein was detected in both cytoplasm and nucleus (Figure 1D). To test if peroxidase activity of APEX2 is maintained in the RPL protein, HEK293T cells were treated with different combinations of BP and H_2_O_2_ 24h post transfection of the RPL plasmid. The detection of biotinylated proteins requires both BP and H_2_O_2_, indicating that APEX2 in the RPL protein retains peroxidase activity (Figure 1E). The results also suggest that endogenous biotinylated proteins are rare in HEK293T cells and low level of endogenous H_2_O_2_ (Belousov et al., 2006; Huang and Sikes, 2014; Lyublinskaya and Antunes, 2019) could not trigger efficient biotinylation. These data suggest that the RPL protein has peroxidase activity and can be applied to target both cytoplasmic and nuclear transcripts.

### Design and validation of gRNAs targeting *U1* snRNA

To test the approach, we asked whether *U1* snRNA proximal proteins (U1 RPPs) identified with the RPL protein include any known *U1* RBPs. The *U1* snRNA was selected for three reasons: (1) its high abundance (Gesteland, 1993), (2) its structures in human *U1* small nuclear ribonucleoprotein (snRNP) and in spliceosome have been solved (Charenton et al., 2019; Pomeranz Krummel et al., 2009; Weber et al., 2010), and (3) its interacting proteins in *U1* snRNP (Stark et al., 2001) and in spliceosome (Zhou et al., 2002) have been well documented.

Since Cas13b targets single-stranded RNA (Cox et al., 2017; Smargon et al., 2017), three gRNAs (*U1*-1, *U1*-2, and *U1*-3) targeting *U1* single-stranded regions were designed based on its structure in pre-B complex (Charenton et al., 2019) (Figure 2A). We first tested if *U1* gRNAs direct wild-type PspCas13b to *U1* and cleave it by measuring *U1* expression in HEK293T cells cotransfected with wild-type PspCas13b plasmid and plasmid expressing *U1* gRNA or nontargeting control (NTC) gRNA at a 1:1 molar ratio. The expression of *U1* was significantly lower in *U1* gRNA-transfected cells compared with NTC gRNA-transfected cells (Figure 2B). The expression of a group of nontargets with a wide range of abundance was not affected (Figure 2B), except *U2*, which may be caused by Cas13b collateral activity (Gootenberg et al., 2018) since *U1* and *U2* are in close contact during spliceosome assembly. The result indicated that *U1* gRNAs can specifically direct PspCas13b to *U1*. We then tested if *U1* gRNAs deliver the RPL protein to *U1* using RNA immunoprecipitation (RIP) experiment. Since the U6 promoter is slightly stronger than CMV promoter in HEK293T cells (Lebbink et al., 2011), a 1:2 molar ratio between the RPL plasmid (CMV promoter) and gRNA expressing plasmid (U6 promoter) was used to avoid nonspecific targeting due to excess RPL protein. The RPL protein was efficiently retrieved by anti-HA but not isotype control IgG (Figure 2C). Analysis of RNA extracted from RIP experiment showed that anti-HA pulled down 5 times more RNA than control (Figure 2D), certifying the RPL protein RNA binding activity. Although there is no significant difference in the amount of RNA pulled down by the RPL protein with NTC or *U1* gRNAs (Figure 2D), *U1* gRNAs significantly enriched *U1* for ∼2-3-fold compared with NTC gRNA (Figure 2E). The fact that much more abundant 18S was not enriched (Figure 2E) suggested that *U1* gRNAs are able to specifically direct the RPL protein to *U1*.

**Figure 2.**
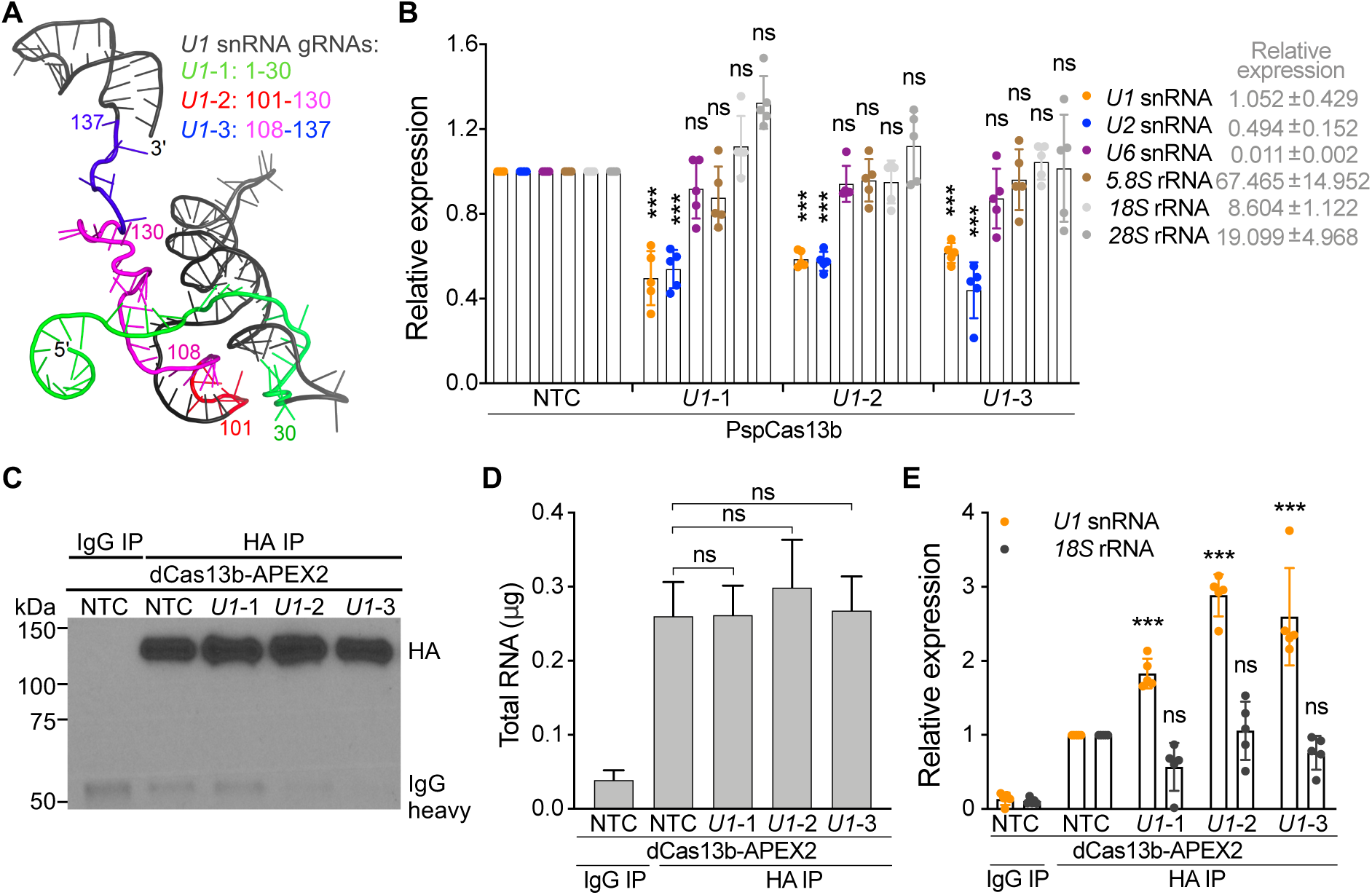
Designing and validating gRNAs targeting *U1* snRNA. **(A).** Based on *U1* structure, three gRNAs with spacers targeting *U1* nucleotides (nt) 1-30 (L/i-1), 101-130 *{U1-2),* and 108-137 *{U1-3)* were designed. Cartoon representation of *U1* (PDB ID: 6QX9, pre-B complex) is colored in black, 1-30 in green, 101-107 in red, 108-130 in magenta, and 131-137 in blue. **(B).** The expression of *U1* snRNA was significantly downregulated in *U1* gRNA-transfected cells. HEK293T cells were cotransfected with plasmid expressing wild-type PspCas13b and plasmid expressing *U1* or NTC gRNA (1:1 molar ratio). The expression of *U1* or a group of nontargets was quantified by RT-qPCR and normalized to *GAPDH.* **(C).** Confirmation of pulldown of the RPL protein by RIP using western blot. HEK293T cells were cotransfected with the RPL plasmid and plasmid expressing *U1* or NTC gRNA (1:2 molar ratio). Anti-HA antibody or isotype control IgG were used to immunoprecipitate the RPL protein. Clean-Blot IP detection reagent was used for blotting. **(D).** The amount of total RNA extracted from RIP experiment. **(E).** *U1* gRNAs specifically directed the RPL protein to *U1.* The expression of *U1* snRNA and nontarget *18S* rRNA was quantified by RT-qPCR and normalized to *GAPDH.* Data shown in **(B), (D),** and **(E)** are mean ± SD from 5 independent experiments. ***p<0.001, ns, not significant. Student’s *t* test.

### RPL-MS identified both *U1* canonical and noncanonical roles-related proteins

We next enriched *U1* RPPs using RPL with the same 1:2 molar ratio to avoid excess RPL protein that can cause nonspecific targeting and proximity labeling. *U1* has compact structure in pre-B complex (Charenton et al., 2019) (Figure 2A) and its size (less than ∼10 nm in diameter) is much smaller than the biotinylating range of APEX2 (likely ∼20-40 nm or larger in diameter) (Fazal et al., 2019; Padrón et al., 2019; Rhee et al., 2013), so we considered experiments using our three *U1* gRNAs as replicates. We analyzed streptavidin-enriched biotinylated proteins by LC-MS/MS (RPL-MS). Using label-free intensity-based absolute quantification (iBAQ) values to measure enrichment in *U1* gRNA relative to protein amounts in the NTC gRNA sample, RPL-MS identified 226 *U1* RPPs (p < 0.05 and log_2_ fold change [FC] > 2, false discovery rate [FDR] < 0.25, Benjamini-Hochberg method), including known *U1* direct RBPs (e.g. SNRNP70, also known as U1-70k) (Stark et al., 2001) and RBPs that likely interact with *U1* indirectly due to their function in the spliceosome (e.g. SNRPA1 and SNRPB2) (Zhou et al., 2002) (Figure 3A, Table S1). We verified the enrichment of U1-70k using western blot and found that it was enriched ∼2-fold by all three *U1* gRNAs (Figure 3B), consistent with RPL-MS results (Figure 3A). Analysis of KEGG pathways enriched in the group of *U1* RPPs using STRING (Szklarczyk et al., 2019) showed that ‘Spliceosome’ is the most significantly enriched pathway (FDR < 10^-8^) (Figure 3C). Indeed, *U1* RPPs included 99 splicing and related factors (Cvitkovic and Jurica, 2013), 56 proteins previously found by *U1* ChIRP-MS (C. Chu et al., 2015), and 58 proteins revealed by XLIP-MS using anti-U1A and/or anti-U1-70k antibody (So et al., 2019) (Figure 3D). In addition, the binding between *U1* and four *U1* RPPs was further supported by corresponding CLIP-Seq data as shown in ENCORI (Li et al., 2014), including DDX3X that is not known to interact with *U1* in human (Deckert et al., 2006; Tarn and Chang, 2009) (Figure 3E). These results together validated that RPL can efficiently identify most known RBPs for *U1*.

**Figure 3.**
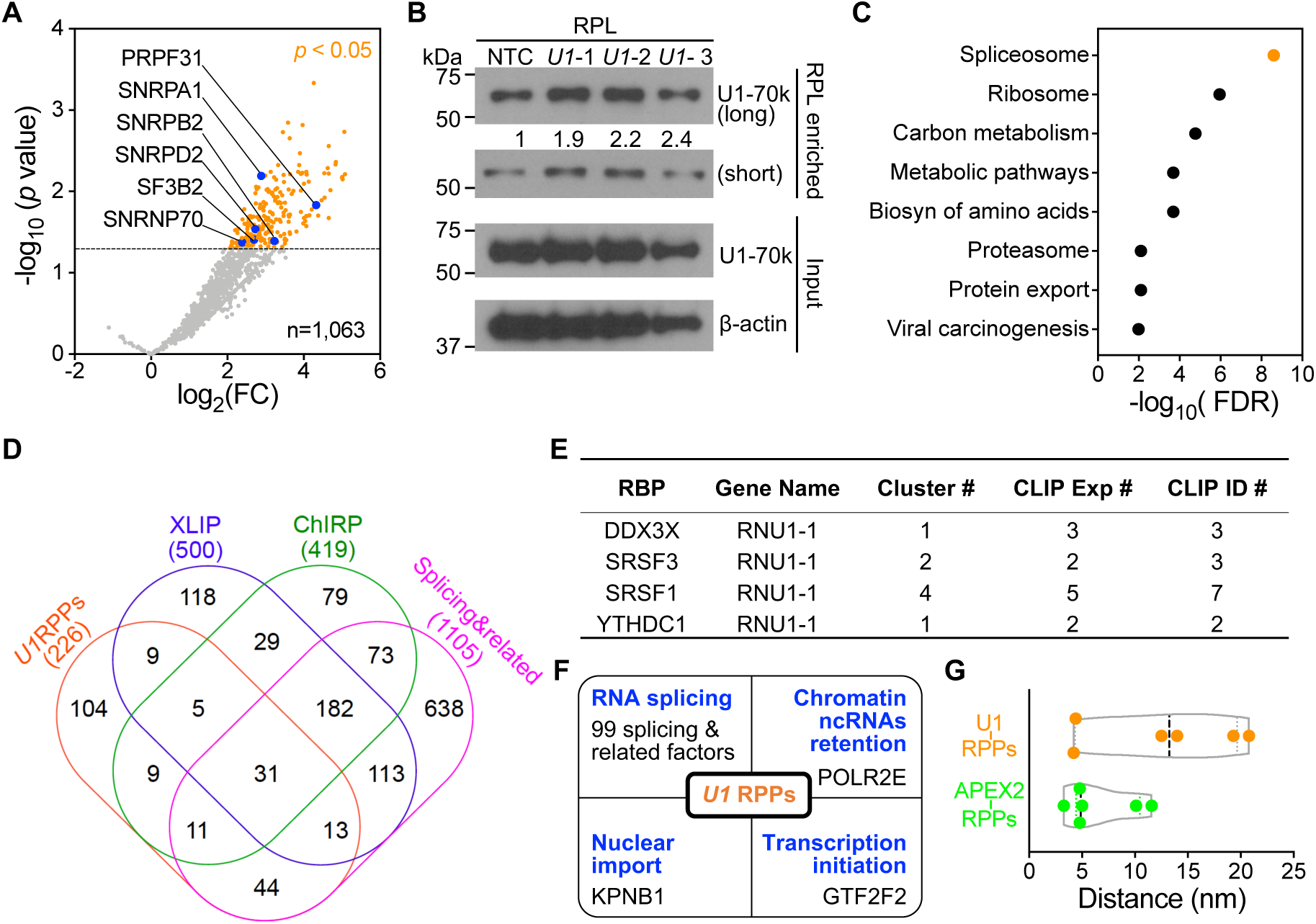
*U1* RPPs identified by RPL-MS. **(A).** *U1* RPPs revealed by RPL-MS include known *U1* RBPs. Volcano plot shows *U1I*NTC iBAQ ratio (fold change, FC) of protein quantification in *U1* gRNA cells compared with NTC gRNA cells. RPL-MS enriched 226 *U1* RPPs (orange dots) that were statistically significant (p < 0.05 and log_2_ FC > 2, FDR < 0.25, Benjamini-Hochberg method). Each dot represents the average value from experiments using three *U1* gRNAs. Blue dots represent proteins from pre-B spliceosome complex. **(B).** *U1* direct RBP U1-70k was enriched by RPL. HEK293T cells transfected with the RPL plasmid and plasmid expressing *U1* gRNA or NTC gRNA were treated with BP and H_2_O_2_. Whole cell lysates (Input) or streptavidin-enriched biotinylated proteins (RPL enriched) were blotted. Numbers represent relative amount of U1-70k under the corresponding conditions from RPL enriched normalized to input. **(C).** KEGG pathways significantly enriched by 226 *U1* RPPs using STRING. **(D).** Comparison of *U1* RPPs, *U1* interactors identified by ChIRP-MS, *U1* interactors identified by XLIP-MS using anti-U1 A and/or anti-U1-70k antibody, and splicing & related proteins. Numbers listed below are total number of proteins from each group. **(E).** List of 4 *U1* RPPs with CLIP-Seq data supporting their association of *U1* found in ENCORI. (F). Summary of *U1* RPPs related to *U1* functions. (G). Inferred distances between *U1* or APEX2 in the RPL protein and those 6 *U1* RPPs present in pre-B complex shown in (A).

*U1* RPPs also included previously reported *U1* interactor RNA polymerase II (Spiluttini et al., 2010; Yu and Reed, 2015) (Figure 3F), which is required for a noncanonical role of *U1* in chromatin retention of ncRNAs (Yin et al., 2020). Moreover, *U1* RPL retrieved proteins involved in chromatin remodeling, DNA modification, histone modification, and transcription (Table S1), which could be regulated by chromatin-associated ncRNAs (Huang et al., 2020; Li and Fu, 2019). The presence of GTF2F2 among the *U1* RPPs may relate to a role for *U1* in regulation of transcription initiation (Damgaard et al., 2008; Kwek et al., 2002) (Figure 3F). Interestingly, RPL-MS revealed nuclear import receptor importing (KPNB1) (Figure 3F), which is required for *U1* nuclear import (Palacios, 1997). Six RBPs in pre-B complex (Charenton et al., 2019) were identified as *U1* RPPs (Figure 3A). Their distances to *U1* snRNA in pre-B complex may provide insight to the biotinylating range of the RPL protein. The inferred distances between APEX2 in the RPL protein and those RBPs are all smaller than 12 nm and the average is 6.6 nm (Figure 3G), suggesting that APEX2 may biotinylate proteins within 12 nm. The inferred distances between *U1* and associated RBPs range from 4.2 nm to 20.8 nm with an average of 12.5 nm (Figure 3G), suggesting that RPL can biotinylate proximal proteins within ∼20 nm of target RNA. These data indicated that RPL enables efficient identification of validated RBPs associated with both canonical and noncanonical functions of *U1*.

### RPL-MS recalled expected categories of proteins for poly(A) tails

To further test the generality of RPL method, we applied it to poly(A) tails, which are adenosines added to the 3’ ends of the majority of eukaryotic mRNAs and many lncRNAs in the absence of template (Derrien et al., 2012; Guttman et al., 2009; Tian, 2005; Yang et al., 2011). Poly(A) tails play critical role in mRNA translation and stability (Dreyfus and Régnier, 2002) and their removal triggers mRNA decapping and decay (Muhlrad et al., 1994; Norbury, 2013; Yamashita et al., 2005). Although the 5’ and 3’ ends of pre-translational mRNAs (Metkar et al., 2018) and deadenylating mRNAs (Chen and Shyu, 2011) are distant (Figure 4A, 5’-3’ distance), the physical distances between the two ends of diverse RNAs are incredibly close regardless of their length, type, species, or complexity (Lai et al., 2018; Leija-Martínez et al., 2014) (Figure 4A, 5’-3’ proximity). As oligomers of 30 nt poly(U) are not found at the 3’ of RNA (Chang et al., 2014; Lim et al., 2018) and rarely occur in the human transcriptome, poly(U)-targeting gRNA, or poly(U) gRNA, was used as negative control. The RPL plasmid was cotransfected with plasmid expressing poly(A) or poly(U) gRNA into HEK293T cells at a 1:2 molar ratio and then RPL was performed. Using label-free iBAQ values to measure enrichment in poly(A) gRNA relative to protein amounts in the poly(U) gRNA sample, RPL-MS enriched 786 proteins as poly(A) tail RPPs (Benjamini-Hochberg-adjusted p < 0.05 and log_2_FC > 2) (Figure 4B, Table S2). Poly(A) tail RPPs included seven poly(A) binding proteins, fifteen 3’UTR binding proteins, ten 5’UTR binding proteins, and one cap binding protein (Figure 4B, Table S3), all of which are known to associate with poly(A) tails. Retrieval of proteins from both 5’ and 3’ ends by RPL within a small radius provided additional evidence for poly(A)+ RNA 5’-3’ proximity.

**Figure 4.**
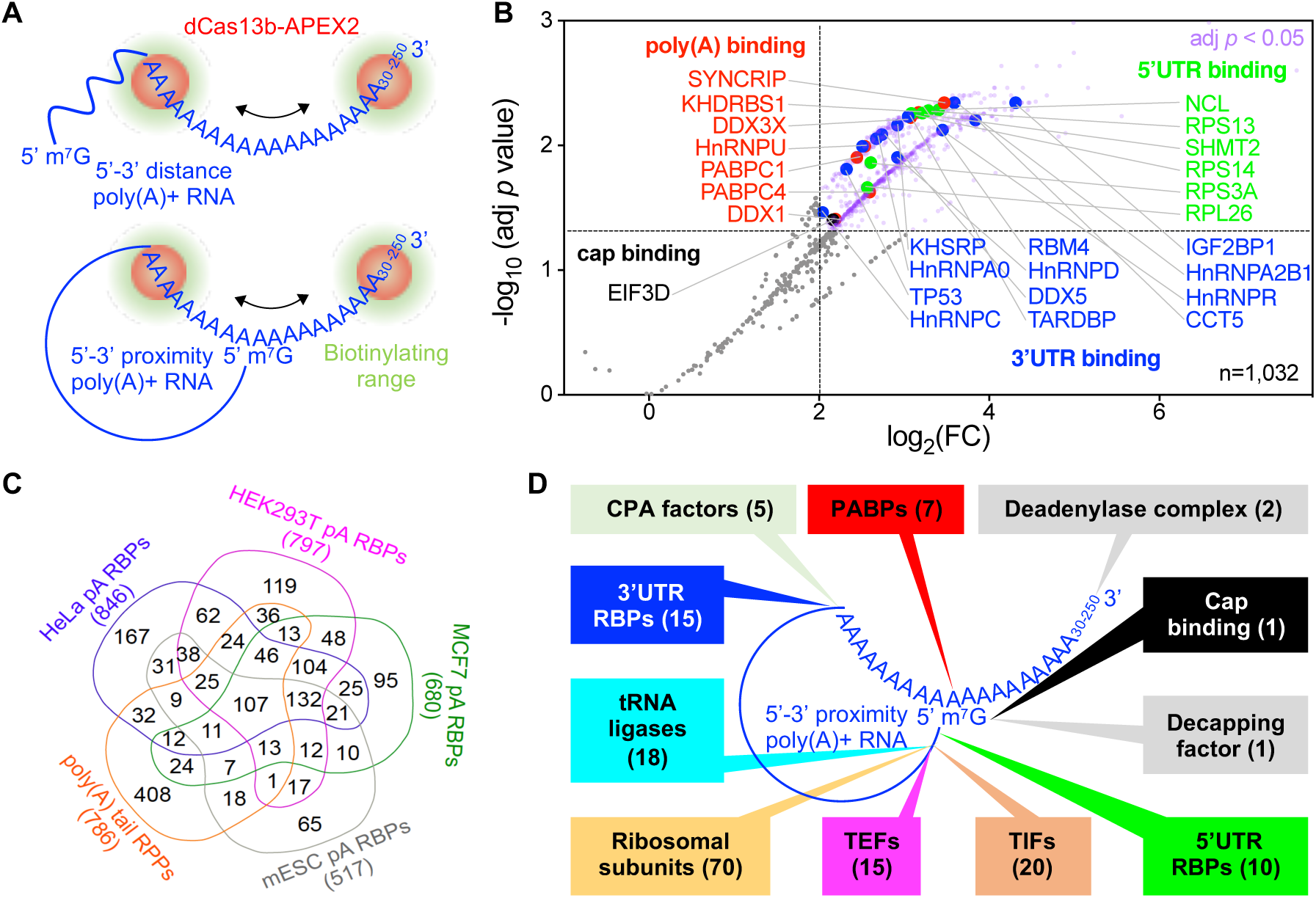
RPL-MS revealed poly(A) tail RPPs in HEK293T cells. **(A).** Application of RPL to poly(A) tails. In the presence of gRNA, the RPL protein (dCas13b-APEX2) is directed to poly(A) tails ranging from 30 nt up to −250 nt. In 5’-3’ distance model, RPL will detect PABPs and 3’UTR binding proteins that bind proximal to poly(A) tail within the biotinylating range. In 5’-3’ proximity model, RPL will also identify cap-binding proteins and 5’UTR binding proteins that bind proximal to the cap and lie within the biotinylating range. **(B).** RPL-MS identified poly(A) tail RPPs. Volcano plot shows RPL-labeled proteins in HEK293T cells. For each protein, the poly(A)/poly(U) iBAQ ratio reflects the enrichment of identified protein in poly(A) gRNA cells compared with poly(U) gRNA transfected cells. RPL-MS identified 786 proteins (light purple dots) as significantly enriched (Benjamini-Hochberg-adjusted *p <* 0.05 and log_2_FC > 2). Each data point represents the average value from biological triplicates. Red dots represent proteins belonging to PABPs, blue dots for 3’UTR binding proteins, green dots for 5’UTR binding proteins, and black dot for cap binding protein. **(C).** Venn diagram shows the comparison of poly(A) tail RPPs and RBPs associated with poly(A)^+^ RNA in different cells. Numbers below each group represent the sizes of the protein cohort. **(D).** Summary of expected categories of poly(A) tail RPPs that support 5’-3’ proximity and the role of poly(A) tail in mRNA translation. Each category of proteins points to a location/region of poly(A)^+^ RNA where they most likely associate with when identified by RPL. PABPs, poly(A) binding proteins; CPA, cleavage and polyadenylation; TIFs, translation initiation factors; TEFs, translation elongation factors.

**Figure 5.**
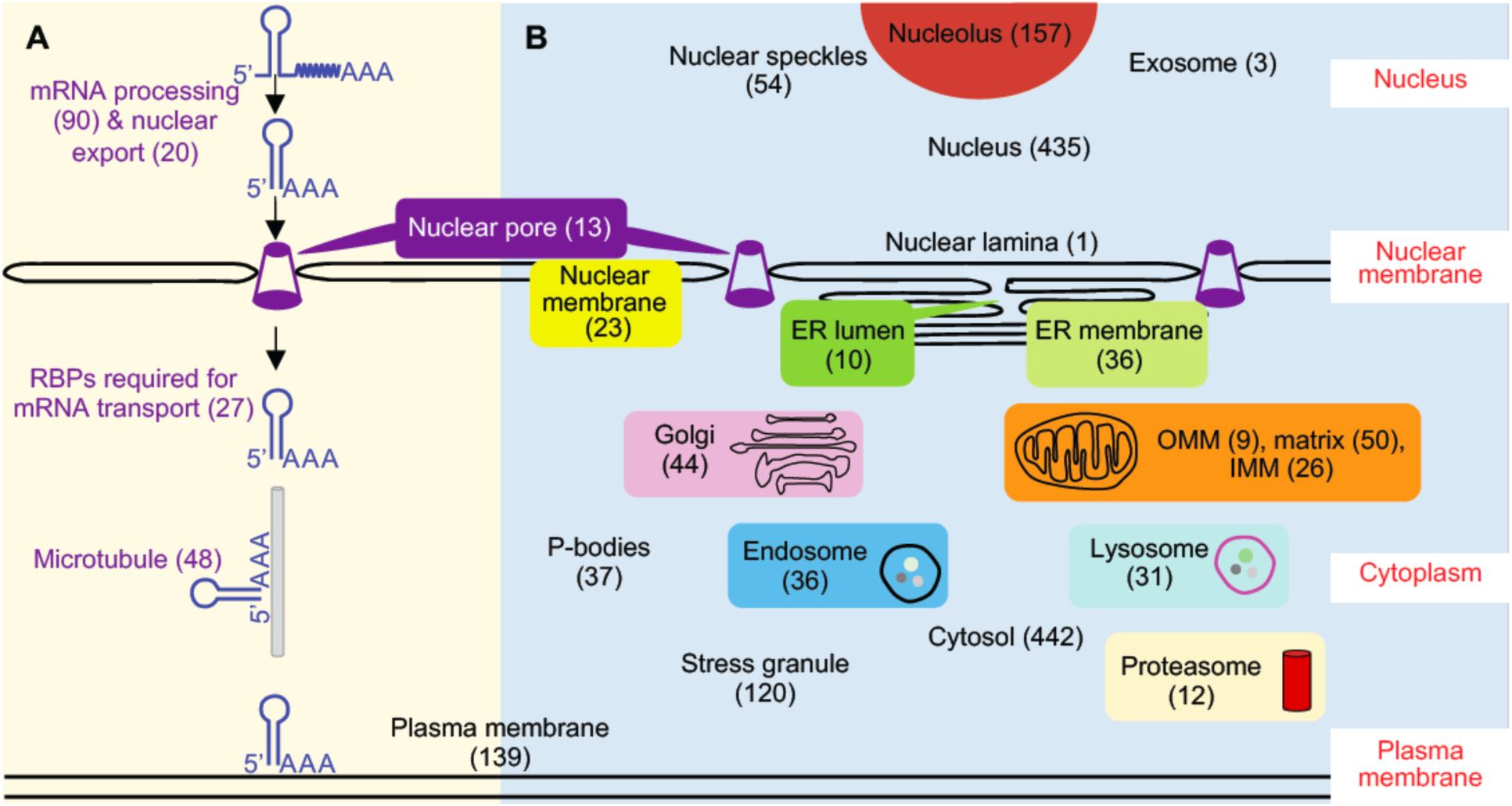
Poly(A) tail RPPs included proteins involved in subcellular localization of RNA. (A). RPL-MS revealed proteins involved in RNA processing, nuclear export, transport, and subcellular localization for poly(A) tail RPPs. (B). A putative subcellular localization map of poly(A)+ RNA built upon subcellular localization of poly(A) tail RPPs. Poly(A) tail RPPs were compared with proteins extracted from corresponding GO terms. Numbers in brackets represent the size of each category of proteins. A full list of proteins in each category can be found in *Table S3*.

Among poly(A) tail RPPs, at least 48% were RBPs interacting with poly(A)+ RNA (Baltz et al., 2012; Castello et al., 2012; Kwon et al., 2013; Milek et al., 2017) (Figure 4C). In *theory, poly(A) gRNA can direct the RPL protein to any transcripts with 30 nt-poly(A) tail or longer* (*Figure 4A*), *including transcripts undergoing polyadenylation, readenylation, deadenylation, or translation. We then interrogated poly(A) tail RPPs for other expected classes of proteins, including factors involved in polyadenylation* (Shi and Manley, 2015), *readenylation or deadenylation* (Yan, 2014), *and translation* (Dreyfus and Regnier, 2002). *Indeed, RPL-MS enriched five cleavage and polyadenylation factors for poly(A)+ RNA* (*Figure 4D*) *but no such factors unique for poly(A)^-^ RNA (e.g. SLBP and ZNF473)* (Gilmartin, 2005) (*Table S3*). *Moreover, poly(A) tail RPPs included three exosome proteins* (Chlebowski et al., 2013), *two deadenylase complex proteins* (Collart, 2016), *as well as decapping factor EDC3* (Mugridge et al., 2018) (*Figure 4D*, *Table S3*). *Importantly, twenty translation initiation factors, fifteen translation elongation factors, seventy ribosomal subunits, and eighteen tRNA ligases were identified by RPL-MS* (*Figure 4D*, *Table S3*), *putatively supporting a model that poly(A) tail recruits translation initiation factors to initiate translation at the 5’ end like their viral counterparts* (Simon and Miller, 2013; Truniger et al., 2017). *Moreover, RPL-MS revealed twelve proteins involved in degradation of AU-rich element-containing mRNAs and 66 nonsense-mediated decay proteins (including 58 ribosomal subunits)* (Chang et al., 2007; Laroia et al., 2002, 1999) (*Table S3*), *further suggesting that RPL enables efficient discovery of most relevant and validated RBPs proximal to poly(A) tails*.

Localization analysis of poly(A) tail RPPs unveils expanded subcellular localizations for poly(A)+ RNA Poly(A) tails are important for RNA nuclear export (Huang and Carmichael, 1996) via the nuclear pore complex (NPC) (Okamura et al., 2015). This is further supported by the presence of 90 mRNA processing factors, 20 mRNA nuclear export proteins, and 13 NPC proteins in poly(A) tail RPPs (Figure 5A). It is not surprising that tRNAs and pre-miRNAs nuclear export factors were also included (Table S3) since their precursors or primary transcripts are also polyadenylated (Cai, 2004; Kadaba et al., 2006). Poly(A) tail RPPs contained eight tRNA processing factors and five tRNA nuclear export factors (Kruse et al., 2000), as well as three pri-miRNA processing factors and two pre-miRNA export factors (Bohnsack et al., 2004; Lund et al., 2004; Yi et al., 2003) (Table S3), supporting that their processing is coupled with export (Kim, 2005; Köhler and Hurt, 2007). Poly(A) tail RPPs recovered 27 RBPs involved in mRNA transport (including zipcodes binding protein IGF2BP1), 48 microtubule proteins, and 139 plasma membrane proteins that are used by mRNAs to achieve different subcellular localizations (Holt and Bullock, 2009; Mofatteh and Bullock, 2017) (Figure 5A), possibly suggesting a role for the poly(A) tail in RNA subcellular localization.

Since unique localizations of RPPs reflect target RNA proximal localizations, we built a putative subcellular localization map for poly(A)+ RNA by comparing poly(A) tail RPPs with proteins extracted from 22 subcellular compartments (Figure 5B, Table S3). The results are generally consistent with previous reports that both mRNAs and ncRNAs have multiple subcellular localizations (Blower, 2013; Carlevaro-Fita and Johnson, 2019; Fazal et al., 2019; Wilk et al., 2016) and also support the presence of mRNAs in P-bodies, stress granule, and the exosome (Chlebowski et al., 2013; Decker and Parker, 2012). Interestingly, RPL-MS also identified marker proteins of the endosome, lysosome, proteasome, and Golgi apparatus, indicative of expanded subcellular localizations for poly(A)+ RNA (Figure 5B, Table S3). Discovery of lysosomal and proteasomal proteins in poly(A) tail RPPs is compatible with the existence of RNA degradation pathway ‘RNautophagy’ in the lysosome (Fujiwara et al., 2013) and degradation function of proteasomes for AU-rich element-containing mRNAs (Laroia et al., 2002, 1999). The identification of endosomal proteins is in accordance with that late endosomes can be used by mRNAs as a platform for translation (Cioni et al., 2019). Poly(A) tail RPPs included Golgi marker c/s-Golgi matrix protein GOLGA2 (Munro, 2011) (Table S3), which has recently been annotated as an RBP by multiple groups (Caudron-Herger et al., 2019; Queiroz et al., 2019; Trendel et al., 2019), suggesting that Golgi may be a novel subcellular location for poly(A)+ RNA. More experimental data are needed to determine which specific transcripts are associated with GOLGA2 in the Golgi apparatus and the biological significance of those interactions.

## Discussion

### RPL: an RNA-centric approach for RPPs identification in living cells

We present an RNA-centric method, RPL, for discovering RPPs for transcripts of interest and evaluate it in two distinct contexts-first interrogating a specific ncRNA target *U1* and second surveying a heterogenous group of poly(A)+ RNA in living cells. Both *U1* RPPs and poly(A) tail RPPs demonstrated that RPL enables efficient discovery of functional relevant RBPs for target transcripts. The recall of KPNB1 for *U1* nuclear import suggests that RPL allows to detect transient and/or weak interacting proteins (Branon et al., 2017; Roux et al., 2013). Compared with alternative methods, RPL needs no crosslinking or sonication, requires far fewer cells (∼20-40 million vs ∼100-800 million) and involves no genetic manipulation, which may interfere target RNA functions (Laprade et al., 2020). The short pulse of labeling potentially permits RPL to be applied to study RNA-protein dynamics. Recently, APEX2 has also been reported to biotinylate proximal nucleic acids (Fazal et al., 2019; Padrón et al., 2019; Y. Zhou et al., 2019), suggesting that RPL could be potentially applied to identify RNA and DNA in addition to proteins proximal to the target RNA (together as ‘RNA proximitome’) within living cells.

During the preparation of our manuscript, similar strategies using different fusion proteins of endonuclease-deficient Cas13 protein (dLwaCas13a, dPspCas13b, and dRfxCas13d) and proximity labeling enzyme (APEX2, BioID2, BASU, and PafA) were reported (Han et al., 2020; Li et al., 2020; Yi et al., 2020; Zhang et al., 2020). Applications of these methods together with ours to both mRNAs and ncRNAs with wide range of abundance (∼10^2^-10^6^ copies/cell) demonstrate that these methods have broad potential to identify functional relevant RBPs for diverse transcripts.

RPPs identified using the dCas13b-APEX2 are expected to include three types of proteins theoretically: proteins that directly bind to target RNA, proteins that associated with target RNA indirectly via protein-protein interactions, and proteins present within the biotinylating range. Biological replicates are expected to help enrich the first two groups of RBPs and reduce the third type as false positive candidates may not be enriched repeatedly. In addition, an optimal molar ratio between the fusion protein and gRNA, which enables efficient proximity-based biotinylation and prevents nonspecific labeling due to excess fusion protein, is crucial for separating signal from noise. A validated set of gRNAs that can specifically direct fusion protein to target RNA with low off-target activity is another key factor. As complementarity between the gRNA spacer and targeted region as well as local RNA accessibility are essential for RNA targeting (Abudayyeh et al., 2017; Cox et al., 2017; Konermann et al., 2018; Smargon et al., 2017), general principles for gRNA designing can provide critical help in choosing spacer sequence and length for gRNA aiming at single-stranded region of target RNA (Bandaru et al., 2020; Wessels et al., 2020).

### Limitations and directions for improvement

High concentration of H_2_O_2_ (1 mM) used in the RPL method may cause oxidative stress and necrosis to the cells (Clément and Pervaiz, 2001) and may preclude the application of RPL to systems sensitive to oxidative stress and cell harm. Similar to other fusion proteins, the RPL protein due to its large size (∼130kDa) may pose steric hindrance to access target RNA and increase the biotinylating range, which could reduce specificity and limit the application to mapping RNA functional domains (Quinn et al., 2014) and RNA-protein interactions at high resolution. Improvement could be achieved using smaller Cas13 proteins by structure-guided truncations (Zhang et al., 2018). Alternatively, CIRTS strategy could be applied to assemble a much smaller gRNA-dependent RNA proximity labeling enzyme (Rauch et al., 2019).

Another limitation is that RPL and similar tools may not identify RBPs for a target RNA as efficiently as antisense probe-based methods, as the RPL protein has to compete with the RBPs bound to the target transcript (Wessels et al., 2020). The RPL protein can only access single-stranded regions of target RNA (Cox et al., 2017; Smargon et al., 2017) and only proteins with electron-rich amino acids like tyrosine exposed on the surface within the biotinylating range have the opportunity to be labeled (Rhee et al., 2013). The same limitation also applies to other proximity labeling enzymes including BioID and its relatives and PafA, which all favor lysine as labeling substrate (Liu et al., 2018; Samavarchi-Tehrani et al., 2020).

### Perspectives

We anticipate that RPL and similar methods will be widely applied to characterize the functions and regulation of diverse categories of RNA in multiple cell types and organisms. Since both Cas13s and proximity labeling are very active research areas, further optimization and refinements of RPL and similar methods are expected. Utilizations of these tools together with protein-centric methods (Licatalosi et al., 2008; Van Nostrand et al., 2016), annotation of RNA structure (Spitale et al., 2015; Sun et al., 2019) could shed light on the molecular mechanisms of lncRNA functions, RNA-protein interactions, RNA functional domain, and binding specificities for RBPs.

## Acknowledgements

This work was supported by the Ovarian Cancer Research Alliance (Ann and Sol Schreiber Mentored Investigator Award 458799 to X.L.) and the National Cancer Institute (K99/R00-CA184415 and R01-CA207456 to K.L.), the Cedars-Sinai Samuel Oschin Comprehensive Cancer Institute (Cancer Biology Grant 231433 to K.L.). We thank Drs. Wei Yang and Bo Zhou at the Cedars-Sinai Medical Center Biomarker Discovery Platform

Core for label-free quantitative mass spectrometry analysis. We also thank anonymous reviewers for their insightful comments and suggestions.

## Author contributions

X.L conceived the project, designed and performed all experiments, and analyzed the data. R.I.C. and M.A.S.F. analyzed the data. K.L. supervised the project. X.L. and K.L. wrote the manuscript with input from all authors.

## Materials and methods

### Key Resources Table

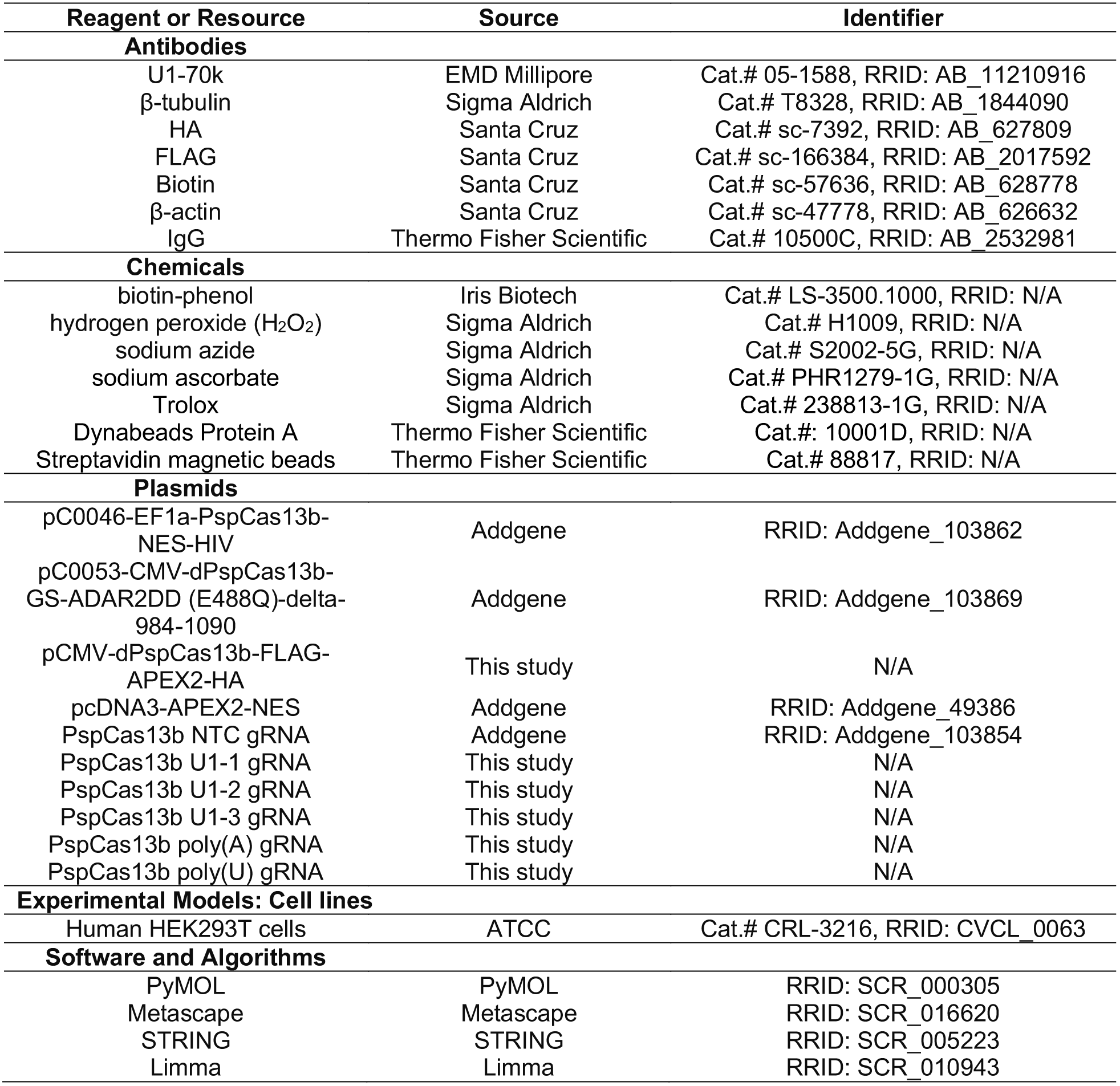

| Reagent or Resource | Source | Identifier |
| --- | --- | --- |
| **Antibodies** |  |  |
| U1-70k | EMD Millipore | Cat.# 05-1588, RRID: AB_11210916 |
| β-tubulin | Sigma Aldrich | Cat.# T8328, RRID: AB_1844090 |
| HA | Santa Cruz | Cat.# sc-7392, RRID: AB_627809 |
| FLAG | Santa Cruz | Cat.# sc-166384, RRID: AB_2017592 |
| Biotin | Santa Cruz | Cat.# sc-57636, RRID: AB_628778 |
| β-actin | Santa Cruz | Cat.# sc-47778, RRID: AB_626632 |
| IgG | Thermo Fisher Scientific | Cat.# 10500C, RRID: AB_2532981 |
| **Chemicals** |  |  |
| biotin-phenol | Iris Biotech | Cat.# LS-3500.1000, RRID: N/A |
| hydrogen peroxide (H<sub>2</sub>O<sub>2</sub>) | Sigma Aldrich | Cat.# H1009, RRID: N/A |
| sodium azide | Sigma Aldrich | Cat.# S2002-5G, RRID: N/A |
| sodium ascorbate | Sigma Aldrich | Cat.# PHR1279-1G, RRID: N/A |
| Trolox | Sigma Aldrich | Cat.# 238813-1G, RRID: N/A |
| Dynabeads Protein A | Thermo Fisher Scientific | Cat.#: 10001D, RRID: N/A |
| Streptavidin magnetic beads | Thermo Fisher Scientific | Cat.# 88817, RRID: N/A |
| **Plasmids** |  |  |
| pC0046-EF1a-PspCas13b-NES-HIV | Addgene | RRID: Addgene_103862 |
| pC0053-CMV-dPspCas13b-GS-ADAR2DD (E488Q)-delta-984-1090 | Addgene | RRID: Addgene_103869 |
| pCMV-dPspCas13b-FLAG-APEX2-HA | This study | N/A |
| pcDNA3-APEX2-NES | Addgene | RRID: Addgene_49386 |
| PspCas13b NTC gRNA | Addgene | RRID: Addgene_103854 |
| PspCas13b U1-1 gRNA | This study | N/A |
| PspCas13b U1-2 gRNA | This study | N/A |
| PspCas13b U1-3 gRNA | This study | N/A |
| PspCas13b poly(A) gRNA | This study | N/A |
| PspCas13b poly(U) gRNA | This study | N/A |
| **Experimental Models: Cell lines** |  |  |
| Human HEK293T cells | ATCC | Cat.# CRL-3216, RRID: CVCL_0063 |
| **Software and Algorithms** |  |  |
| PyMOL | PyMOL | RRID: SCR_000305 |
| Metascape | Metascape | RRID: SCR_016620 |
| STRING | STRING | RRID: SCR_005223 |
| Limma | Limma | RRID: SCR_010943 |

#### Plasmids and cloning

pC0046-EF1a-PspCas13b-NES-HIV was a gift from Dr. Feng Zhang (Addgene plasmid # 103862). pCMV-dPspCas13b-FLAG-APEX2-HA (RPL plasmid) was constructed by replacing ADAR2DD-delta-984-1090 in pC0053-CMV-dPspCas13b-GS-ADAR2DD (E488Q)-delta-984-1090 (a gift from Dr. Feng Zhang, Addgene plasmid # 103869) with FLAG-APEX2-HA subcloned from pcDNA3-APEX2-NES (a gift from Dr. Alice Ting, Addgene plasmid # 49386) using the following primers: dPspCas13b-For: 5’<u>TACCCATACGATGTTCCA</u>GATTACGCT<u>TAAGCGGCCGCTCGAGTC</u>3’, dPspCas13b-Rev: 5’<u>GTCGTCATCCTTGTAGTC</u>GGATCCCAGTGTCAGTCTTTCAAG3’, FLAG-APEX2-HA-For: 5’<u>GACTACAAGGATGACGAC</u>G3’,

FLAG-APEX2-HA-Rev: 5’<u>TGGAACATCGTATGGGTA</u>CTGCAGGGCATCAGCAAAC3’.

PCR was performed using Q5 High-Fidelity DNA Polymerase (New England Biolabs, Cat.# M0491L). PCR fragments were assembled using NEBuilder HiFi DNA Assembly Master Mix (New England Biolabs, Cat.# E2621S) according to manufacturer’s instructions. The following spacer sequences were used to express gRNAs using pC0043-PspCas13b crRNA backbone (a gift from Dr. Feng Zhang, Addgene plasmid # 103854):

NTC: ATGTCTTCCTGGGACGAAGACAA

*U1-1*_1-30_: ATCATGGTATCTCCCCTGCCAGGTAAGTAT,

*U1-2*_101-130_: CAAATT AT G CAGTCGAGTTT CCCACATTT G,

*U1-3*_108-137_: ACTACCACAAATTATGCAGTCGAGTTTCCC,

Poly(A): TTTTTTTTTTTTTTTTTTTTTTTTTTTTTT,

Poly(U): AAAAAAAAAAAAAAAAAAAAAAAAAAAAAA.

The sequences of all constructs have been confirmed using Sanger sequencing.

#### Transfection and *in vivo* proximity dependent biotinylation

For validation of *U1* gRNAs in directing the RPL protein to target *U1,* HEK293T cells were seeded into 12-well plates and were transfected with 1.5 μg the RPL plasmid and 0.5 μg Cas13b gRNAs (NTC, *U1*-1, *U1*-2, *U1*-3) while ∼80% confluency using Lipofectamine 3000 (Thermo Fisher Scientific, Cat.# L3000015). For RIP experiments, HEK293T cells were seeded into 6-well plates and were transfected with 2.5 μg the RPL plasmid and 1.5 μg Cas13b gRNAs while ∼80% confluency using Lipofectamine 3000. For proximity-dependent biotinylation, HEK293T cells were seeded into 150 mm plate and were transfected with 25 μg the RPL plasmid and 15 μg Cas13b gRNAs (NTC, *U1*-1, *U1*-2, *U1*-3, poly[A], poly[U]) while ∼80% confluency using Lipofectamine 3000. HEK293T cells were incubated with 25 mL of DMEM media containing 25 μL of 500 mM biotin-phenol (Iris Biotech, Cat.# LS-3500.1000) in DMSO for 30 min at 37 °C 24h post transfection. Cells were then treated with 1 mM hydrogen peroxide (H_2_O_2_) (Sigma Aldrich, Cat.# H1009) for 1 min on a horizontal shaker at room temperature. The labeling solution was aspirated and cells were washed twice with 25 mL of quencher solution (10 mM sodium azide [Sigma Aldrich, Cat.# S2002-5G], 10 mM sodium ascorbate [Sigma Aldrich, Cat.# PHR1279-1G], and 5 mM Trolox [Sigma Aldrich, Cat.# 238813-1G] in DPBS (Thermo Fisher Scientific, Cat.# 14040182). Cells were then washed three times with 15 mL of DPBS and were pelleted by centrifugation at 1,500 g for 5 min at 4 °C. Cell pellets were snap frozen and stored at −80 °C.

#### Streptavidin enrichment of biotinylated proteins

Cell pellets from two 150 mm plates of transfected HEK293T cells were lysed in 2 mL cell lysis buffer (10 mM HEPES, pH7.5 by KOH, 150 mM NaCl, 0.1% NP-40, 5 mM EGTA, 5 mM Trolox, 10 mM Sodium ascorbate acid, 10 mM Sodium azide, 1 mM PMSF). Streptavidin magnetic beads (Thermo Fisher Scientific, Cat.# 88817) were washed twice with cell lysis buffer and 3.5 mg of each whole cell lysate sample were incubated with 100 μL magnetic bead slurry with rotation for 2 h at room temperature. After enrichment, the flowthrough was removed and beads were washed with 2×1 mL cell lysis buffer, 1mL 1 M KCl, 1 mL 0.1 M Na2CO3, 1 mL of 2 M urea in 10 mM Tris-HCl (pH 8.0), and again with 2×1 mL cell lysis buffer. Biotinylated proteins were then eluted by boiling the magnetic beads in 30 μL 4×Laemmli sample buffer (Bio-Rad, Cat.# 1610747) supplemented with 20 mM DTT and 2 mM biotin.

##### LC-MS/MS and label-free quantitative mass spectrometry proteomic analysis

The streptavidin-enriched proteins were profiled using label-free quantitative mass spectrometry as previously described (B. Zhou et al., 2019) at Cedars-Sinai Medical Center Biomarker Discovery Platform Core.

##### Data analysis for RNA proximal proteins

Data were first filtered to exclude non-human proteins and proteins that were detected in only one or none of the *U1* replicates or poly(A) replicates. Then proteins detected with two or greater unique peptides were subjected to log_2_ transformation. Only the top gene name was kept from multiple candidates. Since *U1* has compact structure in pre-B complex and its size is much smaller than the biotinylating range of APEX2, experiments using *U1* gRNAs (U1-1, *U1*-2, *U1*-3) were considered as replicates to compare with nontargeting controls (NTC1, partially [65%] targeting UBTF; NTC2, targeting poly[A]; NTC3, targeting poly[U]). Moderated t-test with a paired design was used to compare the log_2_-transformed iBAQ values between *U1* and NTC or between poly(A) and poly(U) using limma package (Smyth, 2004*). p* values were adjust using the Benjamini-Hochberg (BH) method (Benjamini and Hochberg, 1995) for multiple comparisons. Proteins with *p* < 0.05 were considered statistically significant. There are 226 *U1* RPPs with *p* < 0.05, log_2_FC > 2, FDR < 0.25 and 786 poly(A) tail RPPs with BH-adjusted *p* < 0.05, log_2_FC > 2.

##### Comparison RPPs with different gene ontology (GO) terms

Lists of human proteins were retrieved (04/13/2020) from QuickGO (https://www.ebi.ac.uk/QuickGO/) *via* searching corresponding GO terms and selecting *‘Homo sapiens (9606)’* under Taxon, except P-bodies and stress granule, which were both curated using data summarized from Wikipedia (04/24/2020) (https://en.wikipedia.org/wiki/P-bodies, https://en.wikipedia.org/wiki/Stress granule). The venn diagrams were generated using online tools (http://bioinformatics.psb.ugent.be/webtools/Venn/).

#### Cellular fractionation

Cells were fractionated as previously described with slight modification (Lin et al., 2019). Six million HEK293T cells were treated with PML buffer (10 mM Tris-HCl, pH 7.5, 0.15% NP-40, 150 mM NaCl) on ice for 4 min after homogenization by flicking. Lysates were loaded onto a 24% sucrose cushion (24% RNase-free sucrose in PML buffer) using large orifice tips, and centrifuged at 15,000 × *g* for 10 min at 4°C. The supernatant (cytoplasmic fraction) was retained and the pellet (nuclear fraction) was washed with 1 ×PBS/1 mM EDTA and resuspended in 200 μL of 1×PBS/1 mM EDTA. Fractionation efficiency was validated by western blot using β-tubulin (Sigma Aldrich, Cat.# T8328, 1:2,000) as cytoplasmic marker and *U1*-70k (EMD Millipore, Cat.# 05-1588, 1:1,000) as nuclear marker.

#### RNA Immunoprecipitation (RIP)

RIP was performed as previously described with slight modification (Lin et al., 2019). Twelve microliter Dynabeads Protein A (Thermo Fisher Scientific, Cat.#: 10001D) were washed with 200 μL HBS (150 mM NaCl, 10 mM HEPES, pH7.5 by KOH) and incubated with 2 μg anti-HA (Santa Cruz, Cat.# sc-7392) or 2 μg rabbit IgG isotype (Thermo Fisher Scientific, Cat.# 10500C) in the presence of 80 μL HBS buffer at room temperature for 1 h. Eight million HEK293T cells were lysed with 800 μL cell lysis buffer (HBS, 0.1% NP-40, 5 mM EGTA, supplemented with 1 χ protease inhibitor cocktail [Roche, Cat.# 11873580001], 1 χ PhosSTOP protease inhibitor cocktail [Roche, Cat.# 4906837001], 1 mM PMSF [Sigma Aldrich, Cat.# 93482], and Superase-in [Ambion, Cat.# AM2696]) at 4°C for 1 h. Cell debris and insoluble proteins were removed by centrifugation at 4°C, 12,000 *g* for 10 min, and the supernatants were incubated with HA-conjugated or IgG-conjugated Dynabeads at 4°C for 1 h. The Dynabeads were then washed 3 times with wash buffer (HBS, 0.1% NP-40) and aliquoted into two halves. Proteins associated with half of the Dynabeads were eluted with 22 μL 4 χ Laemmli sample buffer (Bio-Rad, Cat.# 1610747) by boiling at 95 °C for 5 min. RNA was extracted from the other half of Dynabeads using TRIzol LS (Thermo Fisher Scientific, Cat.# 10296028).

#### RNA extraction and RT-qPCR

RNA associated with immunoprecipitated RPL fusion protein or RNA from gRNA transfected cells were extracted using TRIzol LS. M-MLV reverse transcriptase (Promega, Cat.# M5301) and random hexamers (Promega, Cat.# C1181) were used for reverse transcription. Gene expression was quantified by RT-qPCR using iQ SYBR Green supermix (Bio-Rad, Cat.# 170-8886). The relative gene expression was calculated using the 2^-ΔΔα^ method and normalized to GAPDH. Five nanograms cDNA was used for RT-qPCR analysis on CFX96 Touch Real-Time PCR Detection System (Bio-Rad) using the following primer pairs:

U1-RT-For: 5’CCAGGGCGAGGCTTATCCATT3’, *U1*-RT-Rev: 5’GCAGTCCCCCACTACCACAAAT3’; U2-RT-For: TTCTCGGCCTTTTGGCTAAG; U2-RT-Rev: CTCCCTGCTCCAAAAATCCA;

U6-RT-For: GCTTCGGCAGCACATATACTAAAAT; U6-RT-Rev: CGCTTCACGAATTTGCGTGTCAT; 5.8S-RT-For: GGTGGATCACTCGGCTCGT; 5.8S-RT-Rev: GCAAGTGCGTTCGAAGTGTC;

18S-RT-For: 5’CAGCCACCCGAGATTGAGCA3’, 18S-RT-Rev: 5’TAGTAGCGACGGGCGTGTG3’; 28S-RT-For: CCCAGTGCTCTGAATGTCAA; 28S-RT-Rev: AGTGGGAATCTCGTTCATCC; GAPDH-RT-For: 5’TG CCAAATATGATG ACATCAAGAA3’,

GAPDH-RT-Rev: 5’GGAGTGGGTGTCGCTGTTG3’.

#### Western blot

Protein samples were run on 4-20% gradient precast protein gel (Bio-Rad, Cat.# 456-1096) and transferred onto PVDF membrane (Bio-Rad, Cat.# 1704157). After 1 h blocking, membranes were incubated with anti-FLAG (Santa Cruz, Cat.# sc-166384, 1:1,000), anti-HA (Santa Cruz, Cat.# sc-7392, 1:1,000), anti-biotin (Santa Cruz, Cat.# sc-57636, 1:1,000), or anti-β-actin (Santa Cruz, Cat.# sc-47778, 1:2,000) at 4°C overnight. Membranes were washed three times with Tris-buffered saline containing 0.5% Tween 20 (TBST) before incubating with HRP-conjugated secondary antibody at room temperature for 2 h. Then the membranes were incubated briefly with ECL Western Blotting Substrate (Thermo Fisher Scientific, Cat.#: 32106) after three times wash with TBST. The membranes were exposed to HyBlot Autoradiography Film (Denville Scientific, Cat.#: E3018).

#### Distance calculation

The distances between *U1* snRNA and *U1* RPPs identified by RPL in the pre-B complex structure (PDB ID: 6QX9) were measured using PyMOL (Schrodinger, 2020). We used the distance from *U1* snRNA (nucleotide 1) to the proximal residues of *U1* RPPs to estimate the actual distance (D1). Since there is no structure available for PspCas13b, we used the structure of PbuCas13b (PDB ID: 6DTD) to infer the distance between *U1* RPPs and APEX2 in the RPL protein. Basically, the average distances between gRNA (nucleotide 1, 12, and 23 of spacer) and the C-terminus of PbuCas13b, where the APEX2 was fused to, were measured (D2). The inferred distances between APEX2 and RPPs were then calculated as absolute value of the differences between D1 and D2.

#### Data availability

Raw images for western blots and raw mass spectrometry data for both *U1* RPPs and poly(A) RPPs are included as supporting files.

